# Volatile organic compounds from entomopathogenic and nematophagous fungi, repel banana black weevil (*Cosmopolites sordidus*)

**DOI:** 10.1101/2020.07.03.186429

**Authors:** Ana Lozano-Soria, Ugo Picciotti, Federico Lopez-Moya, Javier Lopez-Cepero, Francesco Porcelli, Luis Vicente Lopez-Llorca

## Abstract

Fungal Volatile Organic Compounds (VOCs) repel banana black weevil (BW), *Cosmopolites sordidus* (Germar, 1824), the key-pest of banana (*Musa* spp.). The entomopathogens *Beauveria bassiana* (Bb1TS11) and *Metarhizium robertsii* (Mr4TS04) were isolated from banana plantation soils using an insect bait. Bb1TS11 and Mr4TS04 were pathogenic to BW adults. Bb1TS11, Bb203 (from infected palm weevils), Mr4TS04 and the nematophagous fungus *Pochonia clamydosporia* (Pc123), were tested for VOCs production. VOCs were identified by Gas Chromatography/Mass Spectrometry - Solid-Phase Micro Extraction (GC/MS-SPME). GC/MS-SPME identified a total of 97 VOCs in all strains tested. Seven VOCs (C1 – C7) were selected for their abundance or previous record as insect repellents. BW starved adults in the dark showed the highest mobility to banana corm in olfactometry bioassays. C7, produced by all fungal strains, is the best BW repellent (p<0.05), followed by C5. The rest of VOCs have a milder repellency to BW. C1 and C2 (known to repel palm weevil) block the attraction of banana corm and BW pheromone to BW adults in bioassays. Therefore, VOCs from biocontrol fungi can be used in future studies for biomanagement of BW in the field.

## 1. Introduction

Banana black weevil (BW), *Cosmopolites sordidus* (Coleoptera: Curculionidae) is the key pest of banana plant crops [1-4]. BW causes more damage to banana crops than any other arthropod pest. BW infestations alter nutrient and water uptakes causing a decline in plant vigor, size and toppling [3-5]. *C. sordidus* originates from South-East Asia [6] and has been invading Central Africa, Central America, the Pacific Islands and all regions where bananas are cultivated between 30°N and 31°S latitudes [7]. In Europe, BW is present in Madeira (Portugal) [8] and the Canary Islands (Spain) [9]. *C. sordidus* can cause losses from 30% up to 90% of the total banana crop yield production in intense outbreaks [10-12].

Banana black weevil’s adults are active at night and prefer humid environment thriving in crop residues. They are slow-moving and poor flyers despite its functional wings. Therefore, the pest dispersion is mainly passive occurring through the handling of infested plant material. Females have low fertility, laying 1–4 eggs/week with considerable intervals during the adult life and, reaching 10–270 total eggs in their lifetime [13]. BW larvae go through 5-8 stages and the post-embryonic development is completed within 5-7 weeks under tropical conditions [13]. BW larvae damage the banana plants boring the corm for feeding [14]. BW tunnels reduce nutrient absorption and can weaken the plant. This leads to reductions in fruit production or falling of bunches [13]. In massive infestations, plants can rot and die.

Banana weevils have several predators and pathogens. BW pathogens include entomopathogenic fungi (*Metarhizium anisopliae* and *Beauveria bassiana*) and nematodes. *B. bassiana* [(Balsamo) Vuillemin, 1912] and *M. anisopliae* [(Metschnikoff) Sorokin, 1883] of the Hypocreales (Ascomycota) have been used for biological control of important agricultural pests [15-18]. These fungi have potential for *C. sordidus* biocontrol under laboratory [19] and field [5] conditions.

*Pochonia chlamydosporia* [(Goddard) Zare and Gams, 2001] is a nematophagous fungus very close phyllogenomically to *M. anisopliae* [20, 21]. *P. chlamydosporia* parasitizes nematode eggs and females [22-24], it is also an endophyte [25, 26] and a soil fungus [27, 28]. Therefore, it has been used as a biocontrol agent of cyst and root knot nematodes [29].

Volatile organic compounds (VOCs) are solid and liquid carbon-based substances that enter the gaseous phase by vaporization at 20 °C and 0.01 kPa [30]. Most VOCs are typically lipophilic liquids with high vapor pressure [31]. The emission of volatile organic compounds plays essential ecological and physiological roles for many organisms, such as fungi which release a broad spectrum of VOCs [32, 33]. Fungi produce volatile compounds through their metabolic pathways [34, 35] and, they appear as intermediate and final products from both primary and secondary metabolism [36]. Fungal VOCs belong to different chemical groups such as monoterpenoids, sesquiterpenes, alcohols, aldehydes, aromatic compounds, esters, furans, hydrocarbons, ketones and others [31, 33, 37]. Since VOCs can spread through the atmosphere and the soil, they are ideal semiochemicals [38]. The aggregation pheromone Sordidine, isolated from BW males [39, 40] is used in insect traps as it is attractant for both sexes. BW adults are attracted to (2R,5S)-theaspirane, the active component of senescent banana leaves [41].

VOCs mediate interactions between organisms such as fungal-insect’s interactions [42]. Fungal production of VOCs is highly dynamic. The VOCs profile of a species or strain may vary according to substrate, culture age, nutrient type, temperature and other environmental parameters [43-45]. VOCs profiles of *B. bassiana* and *M. anisopliae* have been correlated with their pathogenic activity [46]. Some fungal VOCs may have insecticidal or repellent activities. *Muscodor* spp. produces nitrosoamide which kills insects [47]. Naphthalene produced by *Muscodor vitigenus* is an insect repellent [48]. Some insects of various orders avoid fungal species of the Hypocreales detecting their VOCs and negatively responding to them [42, 49-51]. In addition, fungal VOCs show neurotoxicity in *Drosophilla melanogaster* [52].

In this work we isolate naturally occurring entomopathogenic fungi from banana soils to study their capability as biocontrol agents of BW. We analyze their VOCs production to find a new approach for BW management. We also study BW behavior with these VOCs searching for a repellent to be applied in the field.

## 2. Materials and Methods

### Fungal isolation from banana plantations soils using insect baits

Soil samples from banana fields in the Canary Islands (Spain) (Table S1) were sampled as in Asensio *et al.* [53]. For each site two kg of soil were collected from three points, one meter apart from each banana rhizome and between them. Soil was collected from 0-20 cm depth and stored at 4°C until used.

We used *Galleria mellonella* (Bichosa and Cebos Ramiro, Spain) larvae as living baits for entomopathogenic fungi (EF) isolation. Soil samples were screened using a one-millimeter sieve and then dried at room temperature. Forty grams of dried soil from each sample were placed in a petri dish. Three petri dishes (replicates) were prepared per soil sample. Ten milliliters of sterile distilled water (SDW) were then added to each petri dish. Eight living larvae (L3-L4) of *G. mellonella* were buried per plate. Plates were sealed with Parafilm and incubated at 25°C for 15 days in the dark. Plates were shaken periodically to favor contact between larvae and soil. Insects recovered from soil were surface sterilized with 4% sodium hypochlorite for a minute. Insects were then rinsed three times in SDW (5 min each) and finally, dried on sterile filter paper and placed in moist chambers for a week at 25°C in the dark (5 larvae per plate). Larvae were then plated on corn meal agar (CMA) with 50 μg/mL penicillin, 50 μg/mL streptomycin, 50 μg/mL rose Bengal and 1 mg/mL Triton X-100 [54]. Fungi on the larvae were isolated on CMA.

### Morphological identification of EF isolates

Fifteen-day-old colonies from fungal isolates were used to prepare micro-cultures [55]. Fungal isolates were inoculated in 1×1 cm fragments of axenic CMA at their ends on sterile slides. CMA fragments were laid on top with a coverslip and placed in moist chambers. In this way, isolates were determined up to genus level using general taxonomic references [27, 56, 57].

### Molecular identification of fungal strains

EF isolates were grown in potato dextrose broth (PDB) for 5 days at 22°C (120 rpm), filtered and dried to obtain mycelia samples, then they were frozen (liquid nitrogen), lyophilized and stored at -20°C until use. For DNA extraction, we used 30 mg of mycelium for the CTAB (acetyl-trimethyl-ammonium bromide) method of O’Donnell *et al.* [58]. DNA quality was assessed with a ND-1000 Nanodrop (Wilmington, USA).

ITS region (internal transcribed spacers) [59, 60] and the TEF1α gen (translation elongation factor 1-alpha) [61] of EF isolates were amplified and sequenced. Primers used were: ITS-1F (5’TCCGTAGGTGAACCTGCGG3’) and ITS-4 (5’TCCTCCGCTTATTGATATGC3’); EFI-728F (5’CATCGAGAAGTTCGAGAAGG3’) and TEF-1R (5’GCCATCCTTGGGAGATACCAGC3’) fungal specific primers (0.5 μM). PCR was performed with the kit TaqPolimerase 2X (VWR) using the following PCR program: 95°C for two minutes for the initial denaturation and, 35 cycles of 60 seconds at 95°C, 30 seconds at 60°C and 45 seconds at 72°C to facilitate polymerization, finally a five minutes extension at 72°C was included. PCR products were purified using GeneJET Gel Extraction Kit (Thermo Scientific, Fisher) and sent for sequencing. We obtained consensus sequences of the amplified regions of the fungal strains, and they were compared by homology with other sequences in the NCBI database. Match sequences with homology values higher than 95% were used to assign the associated species checked in the NCBI database. The results were crosschecked with identifications obtained microscopically.

### Pathogenicity assays

Virulence of EF isolated from banana soils was evaluated by pathogenicity bioassays. We also included in the bioassays *B. bassiana* 203 strain (from naturally infected *Rynchophorous ferrugineus* adults in SE Spain, Daimès, Elche; CBS 121097) [62]. We evaluated EF virulence by testing the survival of *G. mellonella* larvae inoculated with a given strain of EF under laboratory conditions. Fifteen living larvae of *G. mellonella* were inoculated following the method of Ricaño *et al.* [63]. In the CMA plates bioassays, five larvae (L3-L4) were placed in 9 cm Petri dishes with a 24-day colony of each EF isolate grown in CMA. Insects were kept on the plate for 5 min, shaking the plate regularly to favor contact with the fungus. For controls, plates with non-inoculated CMA medium were used. Pathogenicity assays were carried out in duplicate. Insects were then placed in 9 cm Petri dishes in a moist chamber to evaluate the mortality every 12h up to 20 days. In addition to this, dipping bioassays were carried out too, by making a 10 mL solution of 107 conidia/mL and dipping the insects in the solution for five seconds. In these tests we used *G. mellonella* larvae and *C. sordidus* adults. Mortality was scored daily for 20 days.

Mantel-Cox test (p <0.05, log-rank method, survival percentage analysis) was performed in GraphPad Prism 6 (version 6.01, 2012) using insect survival data. Virulence of EF isolates were evaluated by analyzing the evolution of the survival rate of the insects inoculated with EF versus the control treatment. Survival percentage curves were generated, and significant differences were studied and indicated.

### Gas Chromatography - Mass Spectrometry (GC/MS) analysis

Our two more virulent EF strains (*B. bassiana* 1TS11 and *M. robertsii* 4TS04; Table 1) and *B. bassiana* 203 were used for experiments. The nematophagous fungus *Pochonia chlamydosporia* 123 strain (from eggs of *Heterodera avenae* in SW Spain, Seville; ATCC No. MYA-4875; CECT No. 20929) [64] was also included. Fungi were grown in 250 mL flasks with 75 g of autoclaved rice (*Oryza sativa* cv Redondo; commercial animal food) [62] for 10, 20, 30, 40, 50 and 60 days at room temperature. Uninoculated rice was used as control.

**Table 1.** Entomopathogenic fungi isolated from Tenerife Island banana crops soils.

| Fungi spp. | Code | CECT number | NCBI number | Soil Code | Fungal genera | Blastn |  |
| --- | --- | --- | --- | --- | --- | --- | --- |
| | | | | | | ITS | Tef1α |
| *Simplicillium lamellicola* | A.l.6TS01 | CECT 21125 | MK156720.1 | TS01 | *Simplicillium* sp. | *Simplicillium lamellicola* | - |
| *Metarhizium robertsii* | M.r.4TS04 | CECT 21126 | MK156715.1 | TS04 | *Metarhizium* sp. | *Metarhizium robertsii* | *M. robertsii* |
| *Beauveria pseudobassiana* | B.p.19TS04 | CECT 21122 | MK156716.1 | TS04 | *Beauveria* sp. | *Beauveria pseudobassiana* | *B. pseudobassiana* |
| *Simplicillium lamellicola* | A.l.2TS05 | CECT 21123 | MK156718.1 | TS05 | *Simplicillium* sp. | *Simplicillium lamellicola* | - |
| *Simplicillium lamellicola* | A.l.5TS08 | CECT 21124 | MK156719.1 | TS08 | *Simplicillium* sp. | *Simplicillium lamellicola* | - |
| *Beauveria bassiana* | B.b.1TS11 | CECT 21121 | MK156717.1 | TS11 | *Beauveria* sp. | *Beauveria bassiana* | *B. bassiana* |
| *Metarhizium brunneum* | M.b.8TS21 | - | MT703853 | TS21 | *Metarhizium* sp. | *Metarhizium brunneum* | *M. brunneum* |

GC/MS of fungal cultures was carried out using Solid-Phase Micro Extraction (SPME), with a fused silica fiber (1 cm; ∅ 0.110 mm) [65]. Cultures were processed for VOCs identification at 10, 20, 30, 40, 50 and 60 days after inoculation (dai). For each strain, five grams were sampled and placed in a vial (HS, crimb, FB, 20 ml, clr, cert, 100 PK, Agilent Technologies) hermetically capped by a pressure plug with a plastic membrane. Samples were then placed individually in a 60°C thermostatic bath. Absorption of VOCs occurred by exposing the fiber of the holder to the headspace of the vial for 15 min. The holder was inserted into the GC injector (Agilent 5973 Network Mass Spectrometer - Agilent model 6890N gas chromatograph; column: DB624 30 m, 0.25 mm ID 1.4 µm, J&W Scientific) for a 4 min desorption at 150°C in a spitless mode. The chromatography program used had an initial temperature of 35°C for 5 min and a 3°C/min increasing curve until 150°C, then at that temperature for 1 min. Afterwards, a 5°C/min increasing curve until 250°C to finish the analysis (38 min). The ionization source for electronic impact was 70eV at 230°C. A simple quadrupole was used as a detector at 150°C. Wiley275 library was used for identifying VOCs.

After each chromatography run, the software generated a chromatogram and a list of VOCs. Data obtained were processed in order to obtain, for every strain, a list of the Total-VOCs characteristic of each fungus (T-VOCs) which a match ≥ 50 % with the database entries. A second set that includes Major-VOCs (M-VOCs), which a match ≥ 50 % and a peak-height (relative abundance) > 100.000 ppm. Finally, the third set which contains minor-VOCs (m-VOCs) for the compounds that have a match ≥ 50 % and a peak-height between 100.000 and 20.000 ppm.

### Olfactometer bioassays

BW adults were collected from Tenerife (Canary Islands, Spain) using pheromone traps and kept in plastic boxes (40 cm × 30 cm × 21 cm) at 28 ± 0.5°C in the dark. The plastic boxes contained a small container with distilled water, to maintain a moist environment. Healthy BW adults collected from the field were used for bioassays. Fresh banana corm/pseudocorm (*Musa* sp.) pieces (ca. 16g/each) were used daily for olfactometer bioassays.

A two-way olfactometer (Figure S1) was used to evaluate the behavior of BW subjected to olfactory stimuli. At the end of the arms, two “odor chambers” [66] are placed, consisting of two glass tubes into which insert the olfactory stimuli to test. Tests were conducted by placing a single BW adult at the center of the straight arm. All insects tested had 10 min to move and make a choice or stay in the same place. After each test, the BW was stored. The olfactometer was rinsed after each test with ethanol, n-hexane and distilled water and dried with paper towels. Every test consisted of six replicates of 20 individuals each, which were reused every other time.

### Environmental Conditions

In these tests, two conditions were analyzed, in which the attractive activity of the corm/pseudocorm was tested. We placed the natural attractant in one arm of the olfactometer, while nothing was placed in the other arm. We wanted to evaluate the effect of light (L) or darkness (D) on BW behavior and movement as BW mainly displays nocturnal habits [13]. Furthermore, starvation (S) was also tested. One population included BW fed *ad libitum* (No S), the other had BWs starved for at least a week (S). We evaluate the conditions in combination as follows: L-S, L-No S, D-S and D-No S. In this way, we can deduce in which conditions *C. sordidus* has the highest rate of movement for further bioassays.

### Fungal VOCs repellency

In this group, nine bioassays were conducted with starved BW in the dark (D-S). Seven fungal VOCs (C1-C7) (Table S2) and two BW repellents used in commercial banana plantations [67] (garlic, G. and colloidal Sulphur, S.) were analyzed. Styrene (C1) and benzothiazole (C2) were selected because they repel *R. ferrugineus* [68]. *B. bassiana* 203 and 1TS11 both produced C1 and C2. Camphor (C3) and borneol (C4) were chosen for their repellence to insects [69-71]. C4 is produced by the *B. bassiana* strains and C3 only by *B. bassiana* 1TS11. We selected 1,3-dimethoxy-benzene (C5) and 1-octen-3-ol (C6) from *M. robertsii* and *P. chlamydosporia* because of their high abundance (M-VOCs). Finally, 3-cyclohepten-1-one (C7) was present in all fungal strains studied and in high abundance. Sample of all pure compounds were obtained from SIGMA-ALDRICH.

For the tests, in one arm of the olfactometer a piece of fresh corm/pseudocorm was placed, on the other arm 0.5 mL (C1, C2, C5, C6 and C7) or 0.5 g (C3 and C4) of the pure compounds were inoculated by placing them in a miracloth (Merck KGaA) envelop (3.5 cm × 2.5 cm) with 2 g of silica gel (60A; 70-200μ, Carlo Erba). The volume or weight of the compound was added directly to the silica gel. For the fresh garlic slices (local supermarket) and the colloidal Sulphur (SIPCAM JARDÍN S.L.) 15.6 g of each substance were used.

### Pheromone and fungal VOCs attractiveness

In these tests, four bioassays were carried out under D-S conditions to evaluate a sordidine-based pheromone (ECOSordidine30; ECOBERTURA, La Laguna, Tenerife, Spain) used in field traps. This aggregation pheromone, isolated from BW males [39, 40] is attractant for both sexes. The pheromone was compared in the olfactometer individually with no stimulus and C1 and C2, as described above. The pheromone concentration used was 1/16 (1.7e-4 ml/cm^2^) of the concentration of the commercial formulation (0.27 ml/cm^2^). Used of commercial pheromone caused BW overstimulation and prevented movement.

Other bioassays were carried out in the same conditions as the previous ones. In this case, in one arm of the olfactometer corm/pseudocorm was placed, whereas, in the other arm it was the pheromone alone or with C1 and C2.

### Olfactometer bioassays data treatment for analysis

BWs could go to the arm containing corm/pseudocorm or pheromone, E1 (Figure S1). Others could go to the arm containing fungal VOCs, pheromone or no-stimulus, E2. Lastly, they could not move, EC. This triple ethological response has been evaluated with empirical mobility indexes, to summarize BWs behavior in the tests and to use them in the chi-square statistical tests.

We assume that a BW population placed in the olfactometer in the absence of stimuli would be uniformly distributed in the three possible choices maintaining a 1:1:1 (n_E1_=n_E2_=n_Ec_) ratio relationship. We also consider that the individuals remaining in the EC either, not respond to the stimulus or, are repelled by the substance tested. Therefore, three mobility rates can be formulated as follows: I_E1_=n_E1_/N, I_E2_=n_E2_/N and I_EC_=n_EC_/N. Where, n_E1_= number of individuals choosing E1, n_E2_= number of individuals choosing E2, n_EC_= number of individuals remaining in EC and N= total number of individuals tested. Finally, to obtain an index that takes into consideration all these relationships between groups of individuals, we have summarized theses rates into one, Index of Movement (IM): IM= (I_E1_+ I_E2_)/I_EC_= (n_E1_+n_E2_)/n_EC._ Where, 0<IM<+∞, with n_EC_≠0. The closer to 0 the Index of Movement value is, the higher the motionless portion of the population is. Each of the six replicates (N=20) originate an IM. ANOVA tests were conducted on all olfactometer bioassays using index data on the statistical software Rstudio.

## 3. Results

### 3.1. Entomopathogenic fungi are present in soil from banana plantations

Soil sampling was performed in the Canary Islands where banana is the major crop. The number of samples collected per island was proportional to the area devoted to banana cultivation. In Tenerife, the largest island, with banana as a major crop, most soil samples (23 out of 48) were taken. From 48 soil samples, we have detected the presence of entomopathogenic fungi (EF) in 6 soil samples (12.5%). All EF strains were isolated from soils of Tenerife (Table 1). All field locations with EF are under Integrated Production certification and drip irrigated (Table S1). We have isolated five fungal species from banana soils: *Beauveria bassiana* (1TS11) and *B. pseudobassiana* (19TS04), *Metarhizium robertsii* (4TS04) and *M. brunneum* (8TS21) and three *Simplicillium lamellicola* strains (2TS05, 5TS08 and 6TS01). We have identified them morphologically and by ITS and TEF1α sequencing. NCBI accession numbers are in Table 1. These isolates are deposited into the Spanish Type Culture Collection (CECT). Phylogenetic analysis confirms our species prediction (Figure S2).

### 3.2. Beauveria spp. from banana soils are more virulent than Metarhizium spp. to C. sordidus

Pathogenicity of EF isolates increase with humidity. *G. mellonella* larvae died four days faster under higher humidity conditions (Figure 1A and Figure S3). We established these conditions to evaluate EF pathogenicity in further experiments.

**Figure 1.**
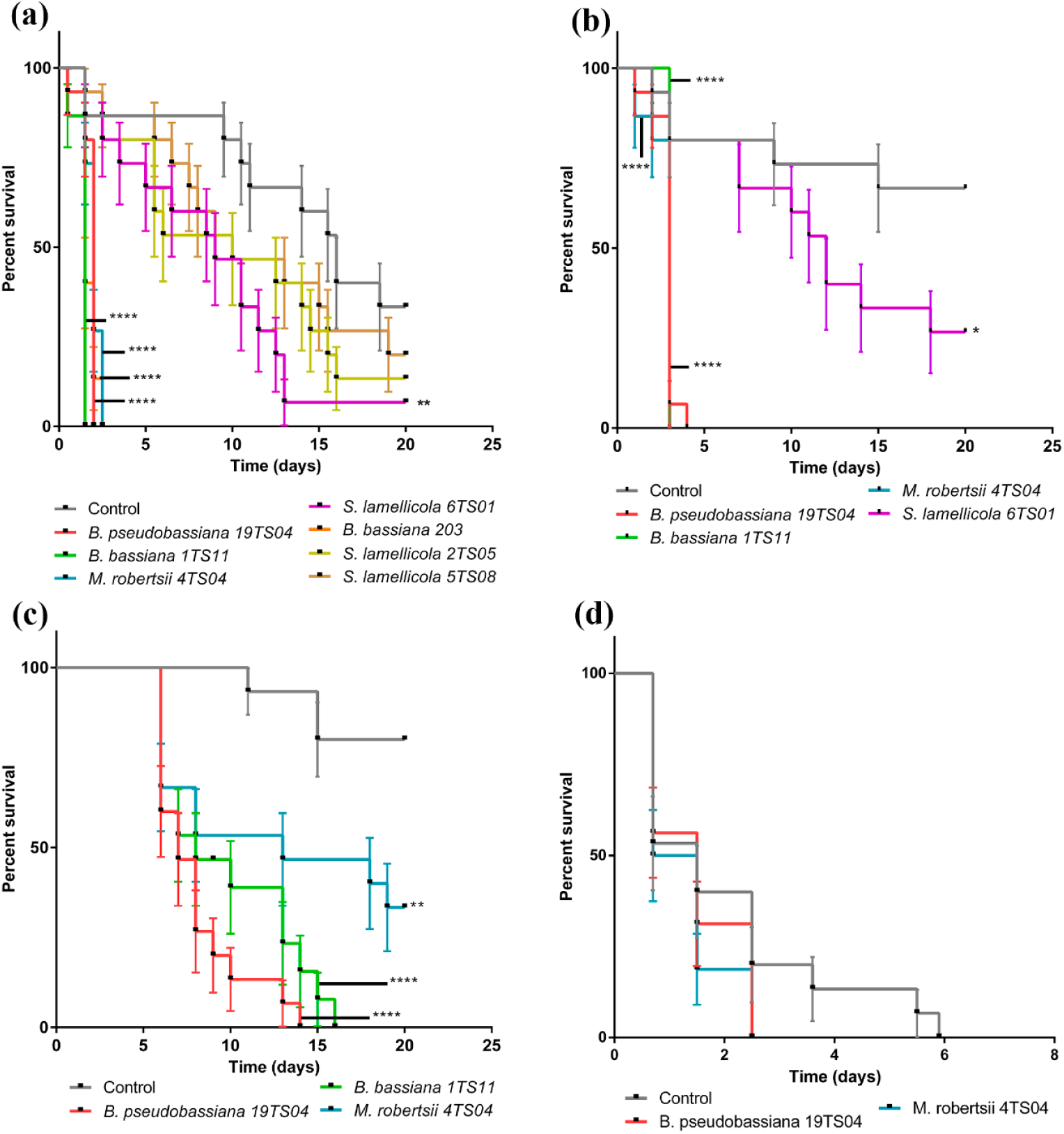
Entomopathogenic fungi isolated from banana crop soils are pathogenic over *Galleria mellonella* larvae and *Cosmopolites sordidus*. (**a**) *Beauveria* spp. and *M. robertsii* isolates are the most virulent on the larvae in the CMA plate bioassay. (**b**) In the dipping bioassay at [10^7^] conidia/mL the most virulent fungi are Beauveria spp. and *M. robertsii*. (**c**) *Beauveria* spp. isolated from banana crop soils of Tenerife are more virulent than *M. robertsii* 4TS04 on *C. sordidus* adults. (**d**) Entomopathogenic fungi isolated from banana crop soils are pathogenic over *Cosmopolites sordidus* larvae. Asterisks indicate significant differences (*p<0.05, **p<0.01, ***p<0.001 and ****p<0.0001) respect to the control.

*Beauveria bassiana* 1TS11 and *B. pseudobassiana* 19TS04 are the most virulent strains, followed by *B. bassiana* 203 and *Metarhizium robertsii* 4TS04. These fungi induced *G. mellonella* full mortality after two days (Figure 1A). Mortality of larvae inoculated with *Simplicillium lamellicola* 2TS05 or 5TS08 do not differ significantly from controls. Therefore, these isolates are not included in further experiments.

In the *G. mellonella* dipping test, the most virulent strain is *M. robertsii* 4TS04, followed by *B. bassiana* 1TS11 and *B. pseudobassiana* 19TS04, with full mortality in less than five days (Figure 1B). So, we have selected these strains to analyze their VOCs.

In the *C. sordidus* pathogenicity test, the most virulent strain is *B. pseudobassiana* 19TS04, followed by *B. bassiana* 1TS11. BW adults are more resistant to EF than *G. mellonella* larvae. This is perhaps because BW cuticle is harder than that of *G. mellonella* larvae (Figure 1C). We also tested BW larvae sensitivity to EF. BW larvae exposed to *B. pseudobassiana* 19TS04 and *M. robertsii* 4TS04 fungal colonies die three days before the uninoculated larvae (Figure 1D). We noticed that BW larvae are very sensitive to manipulation and difficult to rear under laboratory conditions, so we made bioassays with a selection of EF only.

### 3.3. VOCs production by fungal pathogens of invertebrates

To evaluate VOCs, we select the most pathogenic strains (*Beauveria bassiana* 1TS11 and *Metarhizium robertsii* 4TS04) against *G. mellonella* larvae and BW adults. We also study *B. bassiana* 203 isolated from red palm weevil [62] and the nematopathogenous fungus *Pochonia chlamydosporia* 123 [64]. We find 97 volatile organic compounds produced by these fungal strains, during 60 days of growth (Figure 2, Table S3). Only 3-cyclohepten-1-one (C7) and 2-(2-ethoxyethoxy)-ethanol are produced by all fungal strains. *P. chlamydosporia* 123 is the largest VOCs producer with 52 unique compounds, followed by *M. robertsii* 4TS04 (13), *B. bassiana* 1TS11 (8) and *B. bassiana* 203 (5).

**Figure 2.**
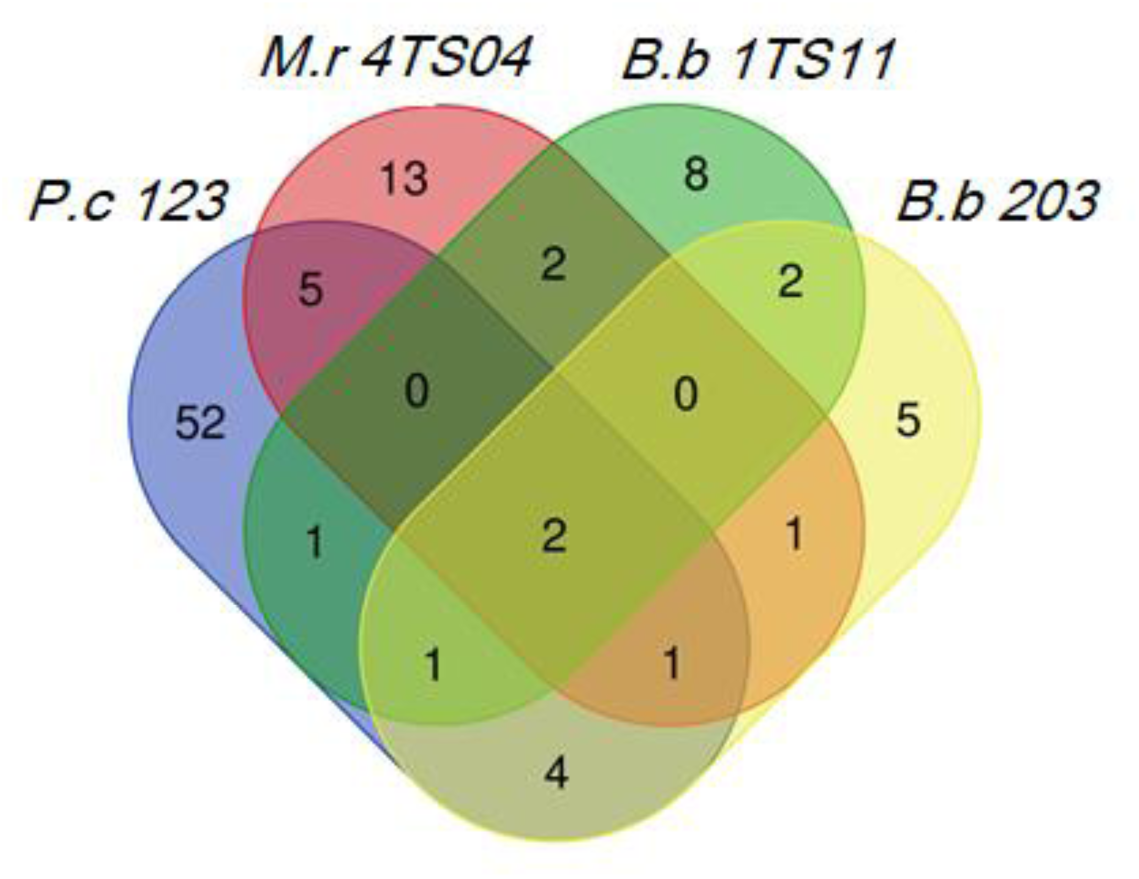
*P. chlamydosporia* 123 is the fungus that produces the most VOCs, followed by *Metarhizium robertsii* 4TS04. Only 2 compounds are produced by all the fungi analyzed by SPME-GC/MS. Abbreviations: *M.r 4TS04* = *Metarhizium robertsii* 4TS04, *B.b 1TS11* = *B. bassiana* 1TS11, *B.b 203* = *B. bassiana* 203 and *P.c 123* = *P. chlamydosporia* 123.

Regarding the VOCs most commonly produced, both *B. bassiana* strains (203 and 1TS11) emit borneol (C4) and 1-octene. In the case of *M. robertsii* 4TS04 and *B. bassiana* 1TS11, they both produce 1,3-octadiene and (+-)-gymnomitrene. Regarding *M. robertsii* 4TS04 and *P. chlamydosporia* 123, they display the highest number of VOCs in common, such as 1-octen-3-ol (C6), 1-methyl-4-(1-methylethyl)-benzene and 1,3-dimethoxy-benzene (C5). *B. bassiana* 1TS11 and *P. chlamydosporia* 123, have only one VOC in common, 1-methylallyl(cyclooctatetraene)titanium. *M. robertsii* 4TS04 and *B. bassiana* 203, have only 4-fluoro-1,2-xylene in common. Finally, both *B. bassiana* 203 and *P. chlamydosporia* 123 emit 2,4-octadiene, β-pinene, α-pinene and 3-octanone.

*Beauveria bassiana* 203 generates 16 compounds characteristic of its metabolic profile (Figure 2). Of these, two are M-VOCs and the other five are m-VOCs (Table 2). 3-cyclohepten-1-one (C7) and borneol (C4) are present in all samplings carried out during the 60 days of growth of the fungus. Most of the compounds detected are found 20 days after inoculation (dai).

**Table 2.** M-VOCs and m-VOCs produced by *Beauveria bassiana* 203 during 60 days of growth. Abbreviations: VOC = volatile organic compound, M-VOCs = major VOCs, m-VOCs = minor VOCs, d = days after inoculation.

| *Beauveria bassiana* 203 | 10d | 20d | 30d | 40d | 50d | 60d |
| --- | --- | --- | --- | --- | --- | --- |
| **M-VOCs** |  |  |  |  |  |  |
| 3-cyclohepten-1-one | X | X | X | X | X | X |
| 4-fluoro-1,2-xylene |  |  |  |  |  | X |
| **m-VOCs** |  |  |  |  |  |  |
| p-Trimethylsilyloxyphenyl-bis(trimethylsilyloxy)ethane |  | X |  |  |  |  |
| Borneol | X | X | X | X | X | X |
| 3-octanone |  | X |  |  |  |  |
| (2R*,6R*,8AS*)-6-hydroxyedulan |  | X |  |  |  |  |
| β-pinene | | | X | | | |

*Beauveria bassiana* 1TS11 also generates 16 VOCs (Figure 2). Two belong to the M-VOCs category, whereas 9 are m-VOCs (Table 3). 3-cyclohepten-1-one (C7) is present in five of the samplings. Borneol (C4) is detected 10, 20, 40 and 50 dai. Benzothiazole (C2) is a m-VOC detected 20 dai only. Most compounds detected are found in the first sampling.

**Table 3.**
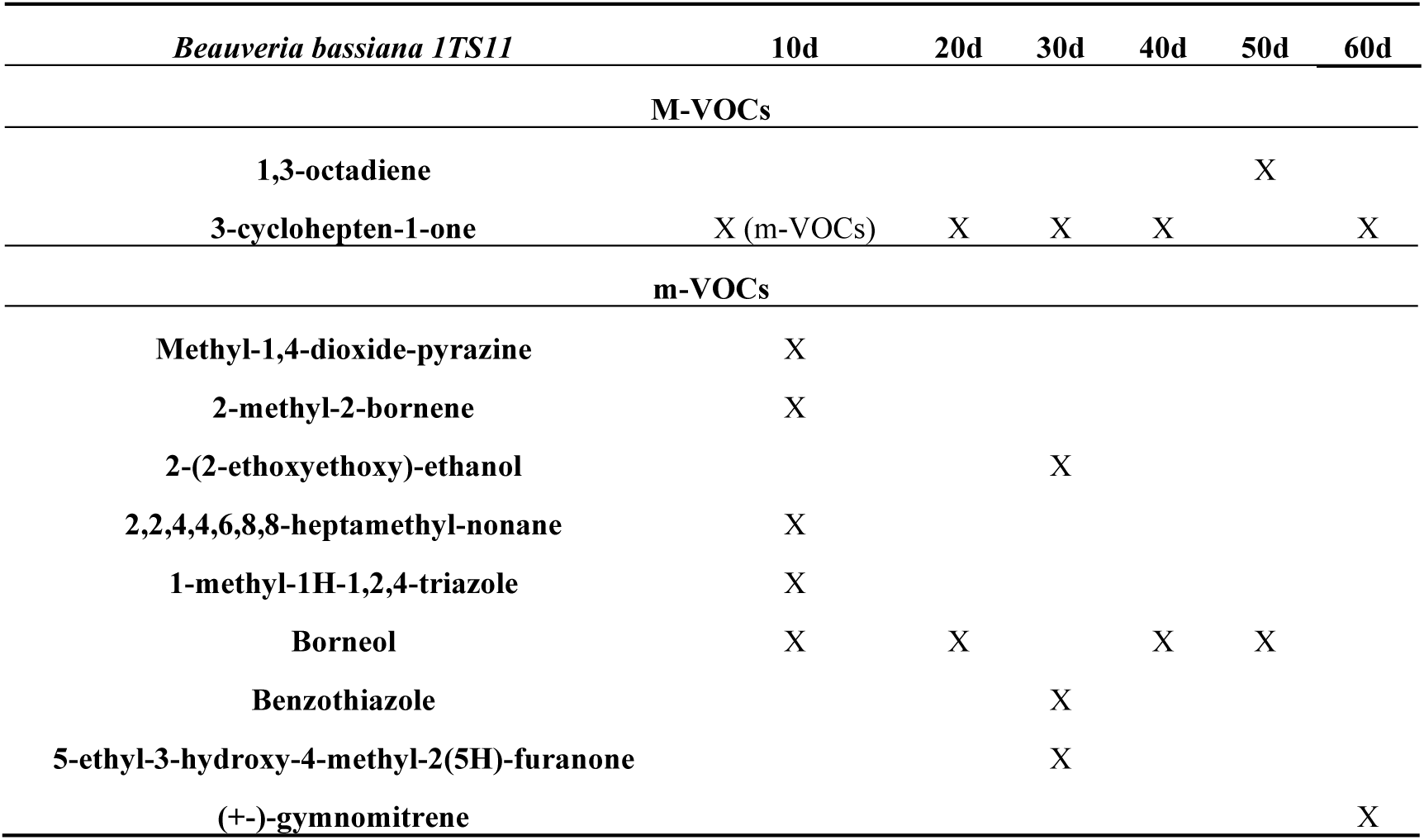
M-VOCs and m-VOCs produced by *Beauveria bassiana* 1TS11 during 60 days of growth. Abbreviations: VOC = volatile organic compound, M-VOCs = major VOCs, m-VOCs = minor VOCs, d = days after inoculation.

*Metarhizium robertsii* 4TS04 shows 24 VOCs in its metabolic profile (Figure 2). Of these, four are M-VOCs and eight are m-VOCs (Table 4). 3-cyclohepten-1-one (C7) is found from 10 to 30 dai, whereas, 1-octen-3-ol (C6) is detected from 10 to 40 dai. 1,3-dimethoxy-benzene (C5) is only found at the first sampling. Most compounds are detected 50 dai.

**Table 4.**
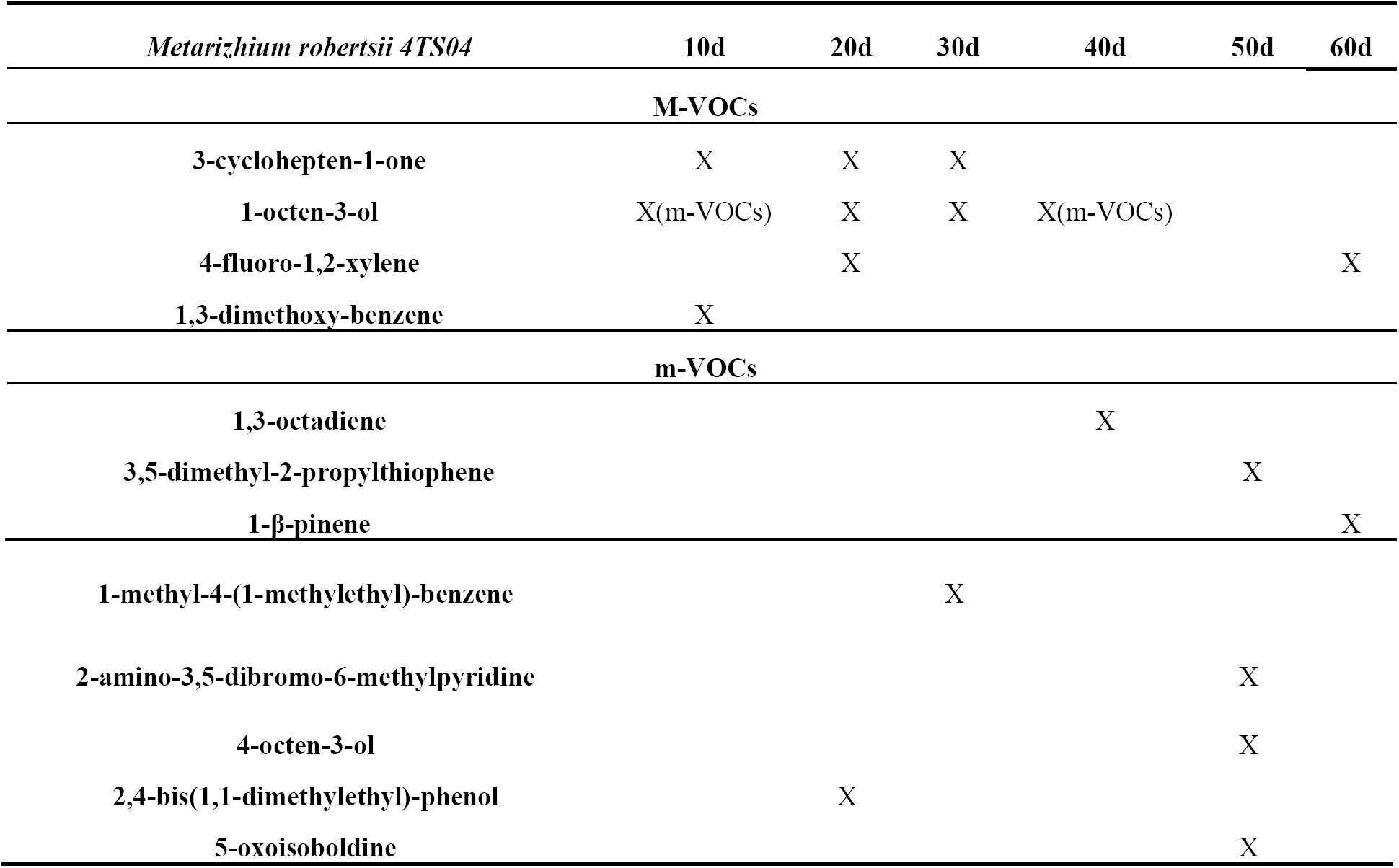
M-VOCs and m-VOCs produced by *Metarizhium robertsii* 4TS04 during 60 days of growth. Abbreviations: VOC = volatile organic compound, M-VOCs = major VOCs, m-VOCs = minor VOCs, d = days after inoculation.

*Pochonia chlamydosporia* 123 shows 66 VOCs (Figure 2). Of these, 12 compounds are M-VOCs and 42 compounds belong to the m-VOCs (Table 5). 1,3-dimethoxy-benzene (C5) is found in the six samplings carried out during the growth of the fungus, while, 1-octen-3-ol (C6) is detected 10, 20, 30 and 60 dai. 3-octanone is present 20, 40 and 50 dai, and 3-cyclohepten-1-one (C7) is found only at the first two samplings. Most compounds identified are detected 40 and 50 dai.

**Table 5.**
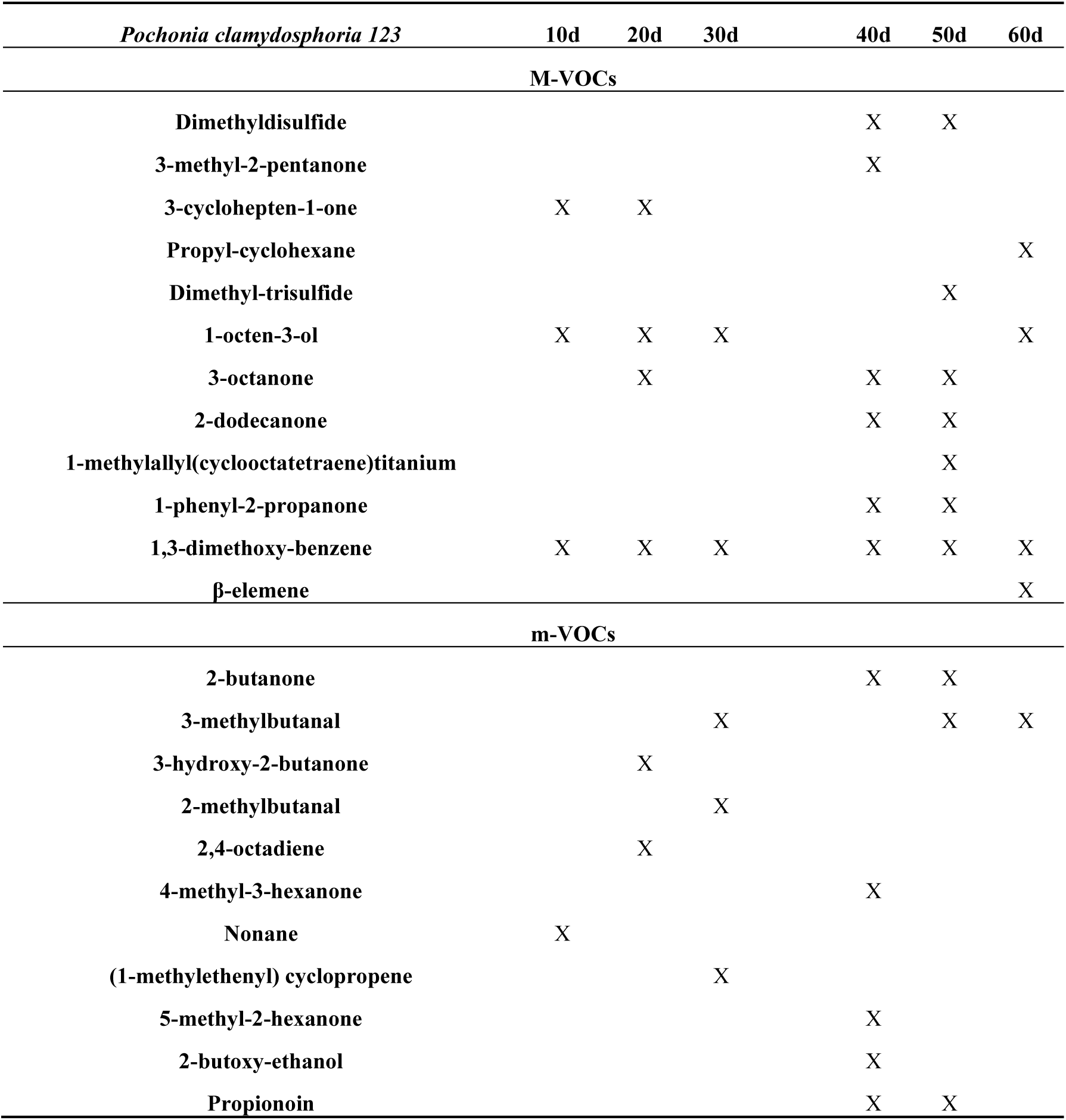
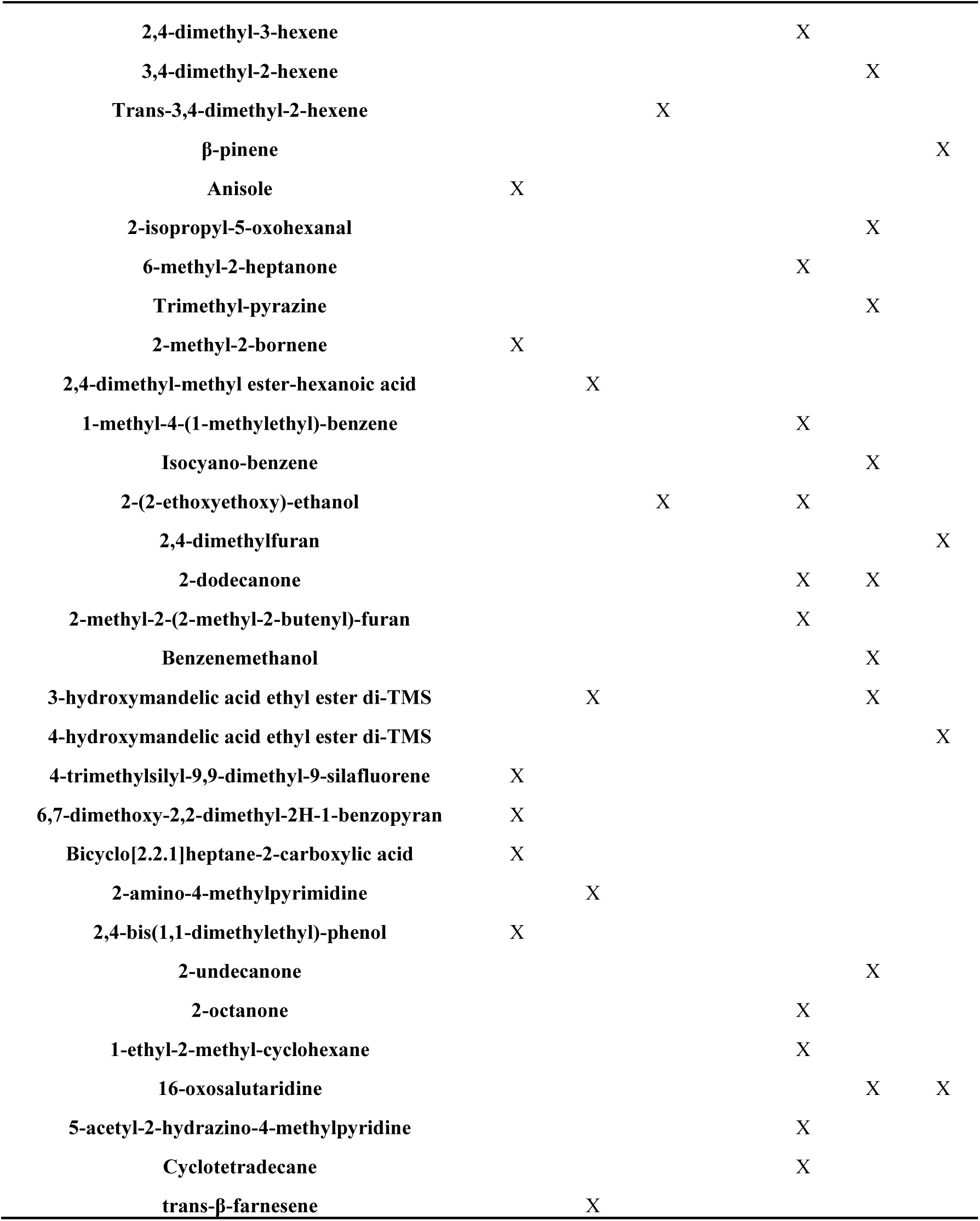
M-VOCs and m-VOCs produced by *Pochonia clamydosphoria* 123 during 60 days of growth. Abbreviations: VOC = volatile organic compound, M-VOCs = major VOCs, m-VOCs = minor VOCs, d = days after inoculation.

### 3.4. Olfactometer bioassays

#### 3.4.1. Effects of environmental conditions on BW mobility

Environmental conditions interfere with BW mobility (Figure 3) (χ^2^ = 17.952; df = 6; p-value = 0.006352; ANOVA: F-value = 3.305; p-value = 0.0413). The nocturnal activity of *C. sordidus* is evident from the higher mobility shown by individuals tested in the darkness, D-S and D-No S (IM_D-S_ = 0.53 and IM_D-No S_ = 0.48) compared to those conducted in the presence of light, L-S and L-No S (IM_L-S_ = 0.36 and IM_L-No S_ = 0.22). Individuals exposed to darkness-starvation conditions (D-S) show the greatest mobility, reflected in the highest Index of Mobility (IM) (IM_D-S_ = 0.53), with significant differences (p<0.05) respect to light-no starvation conditions (L-No S) with the lowest IM (IM_L-No S_ = 0.22). Therefore, BW olfactometer bioassays (next section) are performed under darkness and starvation.

**Figure 3.**
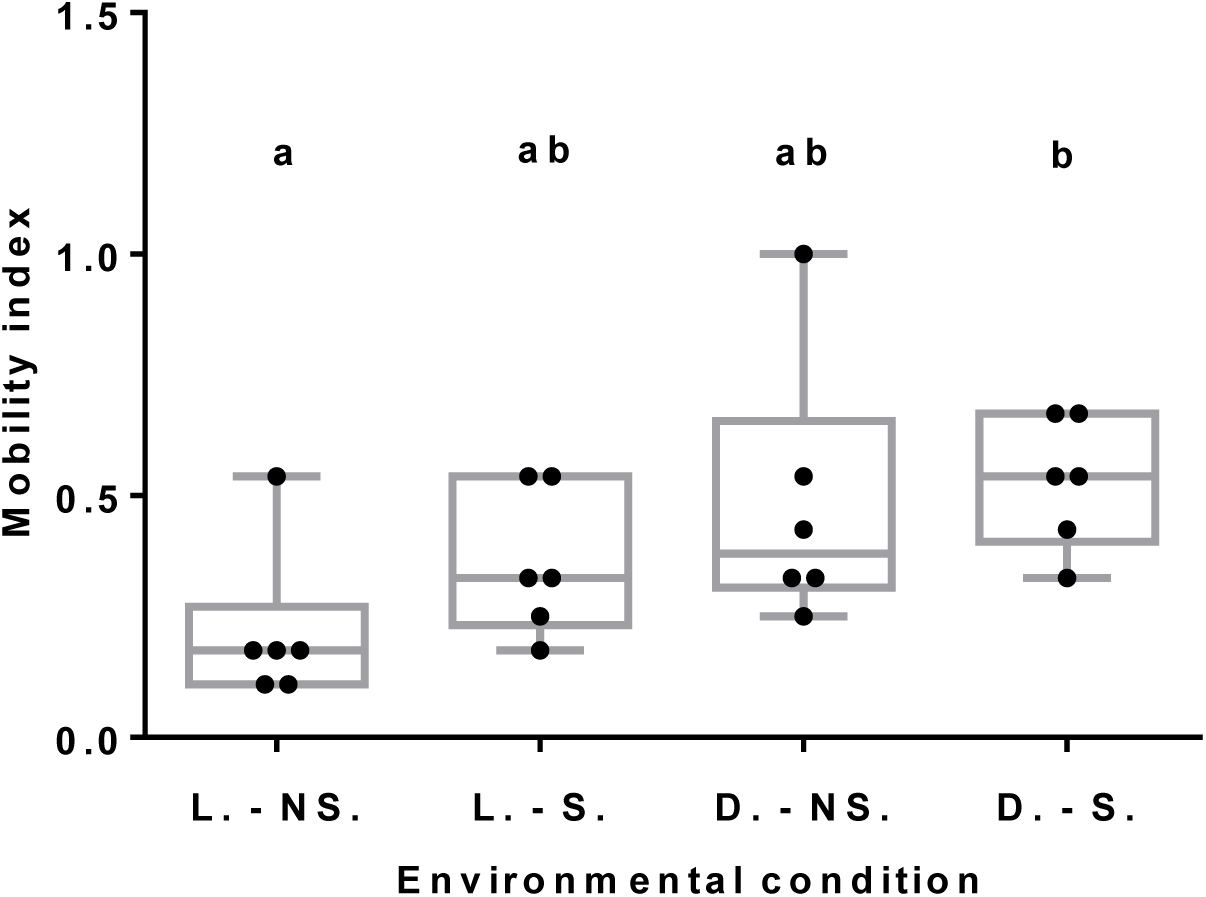
Darkness and starvation are the conditions that generate the most movement on *C. sordidus* (BW), they are the best environmental conditions for analyzing the mobility of BW in olfactometer assays. Abbreviations: L-NS = Light and no starvation, L-S = Light and starvation, D-NS = Darkness and no starvation and D-S = Darkness and starvation. Different letters indicate significant differences between treatments (p<0.05).

#### 3.4.2. Fungal VOCs repel BW

*C. sordidus* mobility is influenced by fungal VOCs (Table S2) and other compounds (technical repellents) tested (Figure 4A) (χ^2^ = 60.881; df = 18; p-value = 1.473e-6; ANOVA: F-value = 3.388; p-value = 0.0026). All fungal VOCs and technical repellents, except for colloidal Sulphur (S.), reduce BW movement compared to the control (corm only, no repellents). However, significant differences (p<0.05) are observed only between the compound 3-cyclohepten-1-one (C7). Sulphur showed the highest IM (IM_S_ = 0.64) compared to the control (IM_D-S_ = 0.53), being significantly different (p<0.05) with 1,3-dimethoxy-benzene (C5) and C7. The compound that mostly reduced BW mobility was C7 (IM_C7_ = 0.11) produced by all fungal strains studied, followed by C5 (IM_C5_ = 0.18) produced by *P. chlamydosphoria* 123 and *M. robertsii* 4TS04 only. Benzothiazole (C2) (IM_C2_ = 0.26) and styrene (C1) (IM_C1_ = 0.28) were the following ones showing a decrease of BW mobility.

**Figure 4.**
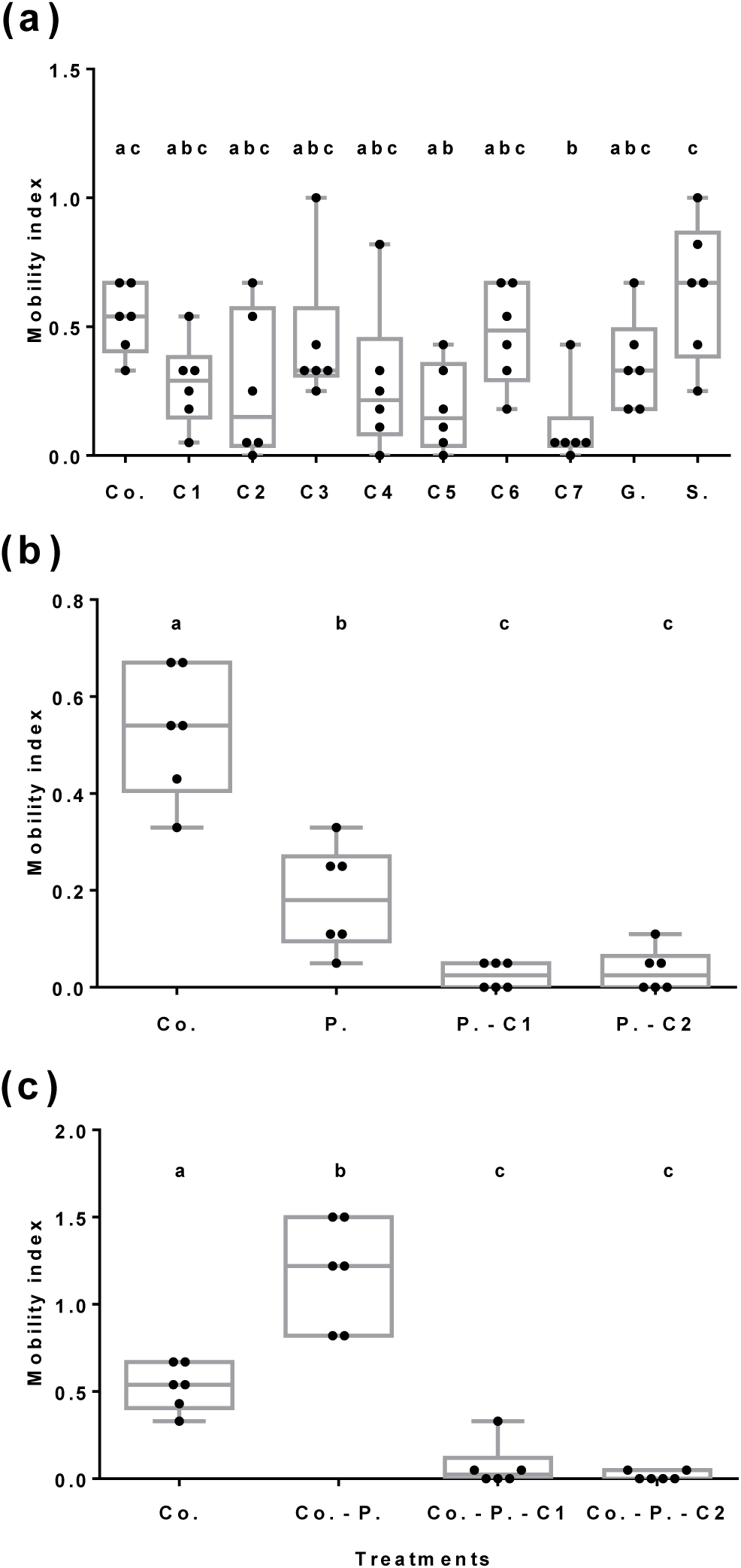
Evaluation of the mobility of *C. sordidus* (BW) subjected to different olfactory stimuli. (**a**) Effect of fungal VOCs and other compounds (technical repellents) versus no stimuli. C7 and C5 are the ones that reduced the most BW mobility. Abbreviations: Co. = Corm, C1 = styrene, C2 = benzothiazole, C3 = camphor, C4 = borneol, C5 = 1,3-dimethoxy-benzene, C6 = 1-octen-3-ol, C7 = 3-cyclohepten-1-one, G = garlic, S = colloidal Sulphur. (**b**) Effect of fungal VOCs on *C. sordidus* pheromone attractiveness. C1 and C2 mask pheromone attractiveness on BW. Co. = Corm vs. no stimuli; P. = Pheromone vs. no stimuli; P.-C1 = Pheromone vs. C1; P.-C2 = Pheromone vs. C2 (**c**) Effect of banana corm, BW aggregation pheromone and fungal VOCs attractiveness to *C. sordidus*. C1 and C2 mask banana corm and pheromone attractiveness on BW. Co. = Corm vs. no stimuli; Co.-P. = Corm vs. Pheromone; Co.-P.-C1 = Corm vs. Pheromone + C1; Co.-P.-C2 = Corm vs. Pheromone + C2. Different letters indicate significant differences between treatments (p<0.05).

#### 3.4.3. Fungal VOCs mask pheromone and banana corm BW attractiveness

*C. sordidus* behavior is influenced by sordidine (BW aggregation pheromone, P), styrene and benzothiazole (C1 and C2, respectively fungal VOCs) (Figure 4B) (χ^2^ = 23.221; df = 4; p-value = 1.14e-4). C1 and C2 significantly modify *C. sordidus* mobility respect to P (ANOVA: F-value = 9.769; p-value = 0.0019). VOCs presence generates an IM (IM_PC1_ = 0.03 and IM_PC2_ = 0.04) lower than that of the pheromone (IM_P_ = 0.18). C1 and C2 therefore repel BW.

Fungal VOCs C1 and C2, combined with pheromone (IM_C-PC1_ = 0.07 and IM_C-PC2_ = 0.02) mask pheromone BW attractiveness. This is reflected by the lower IM of BW of VOCs-pheromone stimuli compared to pheromone alone (IM_C-P_ = 1.18) versus corm (Figure 4C) (χ^2^ = 123.43; df = 4; p-value = 2.2e-16; ANOVA: F-value = 54.13; p-value = <0.0001).

## 4. Discussion

Bananas are essential for food security in tropical and subtropical countries, being one of the best-known, consumed and cultivated fruits [72, 73]. *C. sordidus* is a major pest of banana. It causes more crop destruction than any other arthropod pest in all banana producing countries [5]. BW-resistant banana plants do not cover main commercial cultivars, making this pest a severe problem [74-76].

Biological control agents such as entomopathogenic fungi could be used for BW management [5, 77]. However, this pest has nocturnal habits and passes most of its life cycle within the banana plant and hiding in the leaf litter of the crop, where it is hard to target using EF conidia. As reported in Lopes *et al.* [78], *C. sordidus* adults seem to be less sensitive to EF than many other insects, they performed virulence analysis with indigenous and exotic isolates and observed a lower virulence and recovery rate of the local species.

In this work, we take an alternative approach for BW biomanagement using fungal VOCs. BWs have efficient search mechanisms based on their antennas, specialized primary chemo and mechanoreceptors, which are crucial to ensure survival and reproduction of BWs in the environment [13].

Fungi produce volatile compounds [34,35]. Some of them act as attractants and/or repellents for insects and other invertebrates [42, 46-52]. These compounds may generate alerts in the insect about the presence of possible partners, food, suitable places to lay their eggs or dangers that should be avoided. Therefore, any chemical that could interrupt and modify the behavior of the BW and in general its search ability for the host (*Musa* sp.) could serve as a tool for BW sustainable management.

We have isolated entomopathogenic fungi from banana crop soils in Tenerife (Canary Islands, Spain), in which BW infestations are documented, looking for VOCs repellent to BW. *Beauveria bassiana* (1TS11) and *Metarhizium robertsii* (4TS04) from this survey, both pathogenic to BW, were selected for VOCs analysis. These fungi are common in agricultural fields [79] and banana crops in particular [78]. Since *B. bassiana* 203 produces VOCs repellent to *R. ferrugineus* [68], was included in this study. *P. chlamydosporia* 123, a nematophagous fungus closely related to *M. anisopliae* [20, 21], with a large array of secondary metabolites [80] was also tested for VOCs production. Genomic studies support that some *Metarhizium* species and *P. chlamydosporia* have a single ancestral joint [21].

These fungi produce a total of 97 VOCs. 3-cyclohepten-1-one (C7), a major VOC produced by all fungal strains, was tested on *C. sordidus*. This M-VOC, also produced by other fungi [81], mostly reduces BW mobility among seven VOCs tested. 1,3-dimethoxy-benzene (C5) from *P. chlamydosporia* and *M. robertsii* cultures is the second most repellent VOC to *C. sordidus*.

Camphor (C3) and borneol (C4) produced by *B. bassiana* and *B. pseudobassiana*, show a moderate reduction of BW movement. These are known insects’ repellents [69-71, 82]. 1-octen-3-ol (C6), responsible for the odor of many fungi [83, 84], is a milder repellent to *C. sordidus*.

Styrene (C1) and benzothiazole (C2), are also repellent to *C. sordidus*. They can reduce banana corm and pheromone (sordidine) attractiveness to BW.

## 5. Conclusions

In this work, we have identified new VOCs from EF and a nematophagous fungus, that can be used as BW repellents. 3-cyclohepten-1-one (C7), 1,3-dimethoxy-benzene (C5), styrene (C1), benzothiazole (C2), camphor (C3), borneol (C4) and 1-octen-3-ol (C6) reduce BW mobility and can be, therefore, considered for BW management. In addition, C1 and C2 mask sordidine, a commercial BW aggregation pheromone. Tests should be conducted to determine the effect of selected VOCs as BW repellents in the field. The use of VOCs plus slow-release polymer matrices would improve VOCs performance and durability in the field. The implementation of the technologies associated with the dispersion of these repellents could produce advancement in the agrobiotechnological sustainability of the world banana cultivation. The economic importance of BW at a global level justifies the continuation of research in the identification of new molecules and technologies for the genesis of these new means for bio-management. Managing *C. sordidus*, in an integrated way, could contribute to the increase in banana production, significantly contributing to the increase in global food production, given the extent and importance of this crop.

## 6. Patents

The results regarding fungal VOCs have been patented in the Spanish Office of Brands and Patents (OEPM) with patent number P201930831, with Luis Vicente Lopez-Llorca, Ana Lozano-Soria, Ugo Picciotti, Federico Lopez-Moya and Javier Lopez-Cepero as inventors.

## Author Contributions

Conceptualization, A. L-S, U. P., F. L-M and L.V. L-LL; validation, A. L-S, U. P., F. L-M and L.V. L-LL; formal analysis, A. L-S and U. P.; investigation, A. L-S, U. P. and J. L-C; resources, A. L-S, U. P. and J. L-C; data curation, A. L-S and U. P.; writing—original draft preparation, A. L-S, U. P. and L.V. L-LL; writing— review and editing, A. L-S, U. P., F. L-M, F. P. and L.V. L-LL.; visualization, A. L-S; supervision, F. L-M and L.V. L-LL; project administration, L.V. L-LL; funding acquisition, F. L-M, J. L-C and L.V. L-LL. All authors have read and agreed to the published version of the manuscript.

## Funding

This research was funded by H2020 European Project Microbial Uptakes for Sustainable management of major bananA pests and diseases wit project number 727624.

## Acknowledgments

Authors would like to thank members of the Plant Pathology Laboratory of the University of Alicante, for their help and support. We thank technical support by Lizeth Y. Tabima Cubillos during her stay in the Plan Pathology Laboratory.

## Conflicts of Interest

The authors declare no conflict of interest. The funders had no role in the design of the study; in the collection, analyses, or interpretation of data; in the writing of the manuscript, or in the decision to publish the results.

## Notes

### Competing Interest Statement

The authors have declared no competing interest.

