## Supplementary material for "Volatile organic compounds from entomopathogenic and nematophagous fungi, repel banana black weevil (*Cosmopolites sordidus*)"

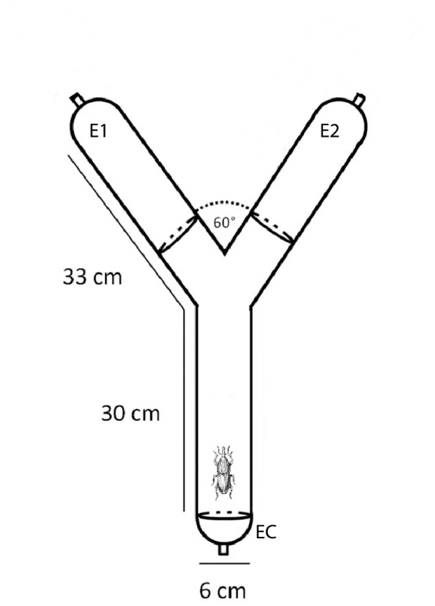

**Figure S1. Two-way olfactometer structure.** BW behaviour in the olfactometer: E1 represents individuals choosing one stimuli, E2 represents individuals choosing the other stimuli or absence of stimulus and EC indicate individuals not moving from the inicial place.

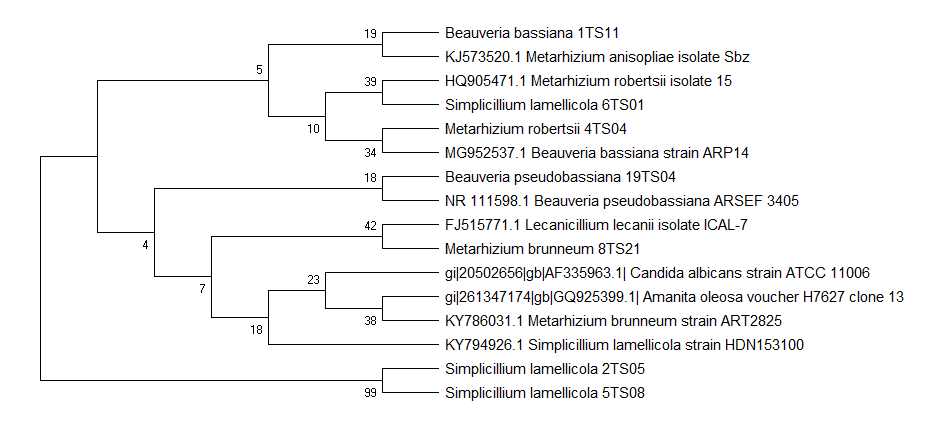

**Figure S2. Molecular Phylogenetic analysis of the ITS region by Maximum Likelihood method (Boostrap consensus tree of 800 replicates), based on the Tamura-Nei model.** Branches corresponding to partitions reproduced in less than 50% bootstrap replicates are collapsed. The percentage of replicate trees in which the associated taxa clustered together in the bootstrap test are shown next to the branches [1]. Evolutionary analyses were conducted in MEGA X [2].

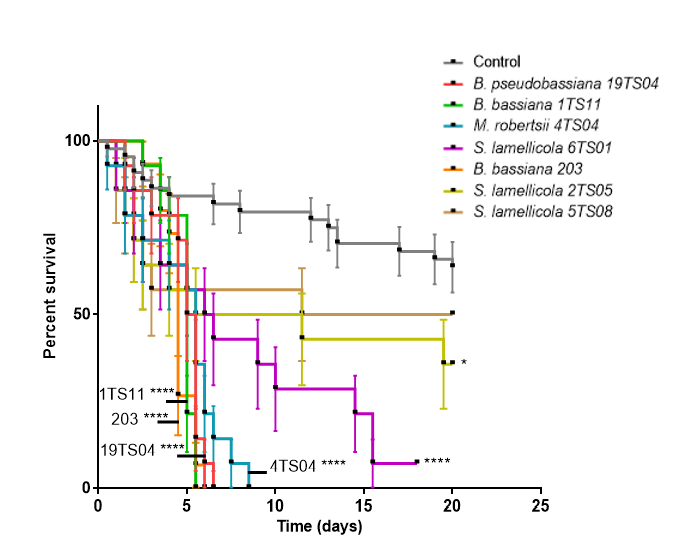

**Figure S3.** *B. pseudobassiana* 19TS04, *B. bassiana* 1TS11, *M. robertsii* 4TS04, *B. bassiana* 203 and *S. lamellicola* 6TS01 are significantly virulent on *G. mellonela* larvae at room humidity conditions. Asterisks indicate significant differences (*p<0.05, **p<0.01, ***p<0.001 and ****p<0.0001) respect to the control.

**Table S1. Soil collection from banana fields in the Canary Islands.** Abbreviations: Lat. = latitude, Long. = longitude, ASL = altitude, S = sprinkling, D = dripping, Cert. = certification, Int. P = integrated production, ECO = ecological crop, Conv. = conventional crop.

| Code | Island | SIGPAC | Lat. | Long. | UTM | ASL | Irrigation | Cert. |
| --- | --- | --- | --- | --- | --- | --- | --- | --- |
| GS01 | La Gomera | 38/21/16/9006/2 | 28°10'24" N | 17°11'17" W | 28 R 285181 3118341 | 82 | S | No |
| GS02 | La Gomera | 38/21/15/13/2 | 28°10'38" N | 17°11'00" W | 28 R 285652 3118764 | 32 | S | No |
| GS03 | La Gomera | 38/21/15/167/1 | 28°10'07" N | 17°11'31" W | 28 R 284790 3117824 | 83 | S | No |
| PS01 | La Palma | 38/14/21/64/1 | 28°30'06'' N | 17°52'16" W | 28 R 218959 3156139 | 56 | S | Int. P |
| PS02 | La Palma | 38/7/6/306/1 | 28°49'49" N | 17°47'00" W | 28 R 228404 3192365 | 170 | S | Conv. |
| PS03 | La Palma | 38/24/13/218/1 | 28°38'26" N | 17°55'09" W | 28 R 214628 3171650 | 267 | S | ECO |
| PS04 | La Palma | 38/24/26/10/1 | 28°34'06" N | 17°53'36" W | 28 R 216961 3163582 | 69 | S | Conv. |
| PS05 | La Palma | 38/24/26/82/1 | 28°33'40" N | 17°53'27" W | 28 R 217186 3162775 | 34 | S | Conv. |
| PS06 | La Palma | 38/14/22/97/4 | 28°29'31" N | 17°52'20" W | 28 R 218824 3155063 | 10 | S | Conv. |
| PS07 | La Palma | 38/30/7/156/1 | 28°44'58" N | 17°44'11" W | 28 R 232781 3183297 | 185 | S | ECO |
| PS08 | La Palma | 38/24/13/56/1 | 28°38'31" N | 17°55'21" W | 28 R 214306 3171812 | 250 | S | Conv. |
| PS09 | La Palma | 38/47/13/560/1 | 28°40'09" N | 17°56'08" W | 28 R 213103 3174861 | 360 | S | Conv. |
| PS10 | La Palma | 38/30/13/252/1 | 28°46'44'' N | 17°45'24'' W | 28 R 230875 3186607 | 70 | S | Conv. |
| PS11 | La Palma | 38/7/6/306/1 | 28°49'49'' N | 17°46'58'' W | 28 R 228459 3192364 | 160 | S | Conv. |
| PS12 | La Palma | 38/14/10/36/2 | 28°29'08'' N | 17°51'59'' W | 28 R 219379 3154341 | 63 | S | Conv. |
| TS01 | Tenerife | 38/26/3/48/ 1 a 6 | 28°24'23" N | 16°30'50" W | 28 R 351703 3143154 | 215 | D | Int. P |
| TS02 | Tenerife | 38/31/9/181/2 | 28°23'28" N | 16°34'18" W | 28 R 346021 3141533 | 230 | D | No |
| TS03 | Tenerife | 38/42/1/32/1 | 28°22'23" N | 16°48'51" W | 28 R 322228 3139866 | 50 | D | Int. P |
| TS04 | Tenerife | 38/18/3/403/2 a 5 | 28°23'41" N | 16°40'12" W | 28 R 336391 3142063 | 115 | D | Int. P |
| TS05 | Tenerife | 38/28/5/9065/9 | 28°24'00" N | 16°33'34" W | 28 R 347231 3142503 | 150 | D | Int. P |
| TS06 | Tenerife | 38/10/5/17/3 | 28°21'53" N | 16°52'32" W | 28 R 316197 3139035 | 60 | D | ECO |
| TS07 | Tenerife | 38/19/8/9000/136 | 28°10'48" N | 16°47'47" W | 28 R 323653 3118448 | 170 | D | ECO |
| TS08 | Tenerife | 38/19/7/34/8 | 28°10'52" N | 16°48'02" W | 28 R 323245 3118577 | 135 | D | Int. P |
| TS09 | Tenerife | 38/1/1/82/1 | 28°08'54" N | 16°47'43" W | 28 R 323710 3114937 | 70 | D | Int. P |
| TS10 | Tenerife | 38/10/51/4 | 28°08'42" N | 16°47'16" W | 28 R 324441 3114557 | 80 | D | Int. P |
| TS11 | Tenerife | 38/6/4/214/6 | 28°01'56" N | 16°38'58" W | 28 R 337858 3101869 | 75 | D | Int. P |
| TS12 | Tenerife | 38/26/1/57/6 | 28°25'04" N | 16°30'43" W | 28 R 351910 3144413 | 100 | D | Conv. |
| TS13 | Tenerife | 38/26/8/43/1 | 28°23'29" N | 16°32'30" W | 28 R 348961 3141526 | 91 | D | Conv. |
| TS14 | Tenerife | 38/26/9/91/2 | 28°23'24" N | 16°33'04" W | 28 R 348033 3141384 | 252 | D | Conv. |
| TS15 | Tenerife | 38/23/1/170/1 | 28°31'51" N | 16°23'47" W | 28 R 363375 3156803 | 75 | D | Conv. |
| TS16 | Tenerife | 38/10/1/146/10 | 28°22'44" N | 16°50'20" W | 28 R 319815 3140550 | 95 | D | Conv. |
| TS17 | Tenerife | 38/42/7/6/16 | 28°22'04'' N | 16°48'13'' W | 28 R 323254 3139266 | 72 | D | ECO |
| TS18 | Tenerife | 38/23/2/44/3 | 28°31'38" N | 16°23'09'' W | 28 R 364403 3156391 | 115 | D | Conv. |
| TS19 | Tenerife | 38/26/1/29/1 | 28°24'36'' N | 16°30'41'' W | 28 R 351953 3143551 | 205 | D | Conv. |
| TS20 | Tenerife | 38/26/2/13/1 | 28°24'38'' N | 16°30'52'' W | 28 R 351655 3143616 | 175 | D | Conv. |
| TS21 | Tenerife | 38/19/5/233/1 | 28°11'36'' N | 16°49'02'' W | 28 R 321629 3119956 | 60 | D | Int. P |
| TS22 | Tenerife | 38/26/3/50/2 | 28°24'37'' N | 16°31'02'' W | 28 R 351382 3143589 | 170 | D | Int. P |
| TS23 | Tenerife | 38/19/2/939/1 | 28°12'51'' N | 16°49'57'' W | 28 R 320164 3122287 | 30 | D | Int. P |
| CS01 | Gran Canaria | 35/09/2/111 | 28°09'28" N | 15°39'01'' W | 28 R 436156 3114851 | 75 | D | Conv. |
| CS02 | Gran Canaria | 35/6/6/1/1 | 28°08'05'' N | 15°31'36'' W | 28 R 448281 3112238 | 145 | D | Conv. |
| CS03 | Gran Canaria | 35/9/2/103/2 | 28°09'27'' N | 15°39'11'' W | 28 R 435883 3114822 | 50 | D | Conv. |
| CS04 | Gran Canaria | 35/6/6/10/1 | 28°08'01'' N | 15°31'14'' W | 28 R 448881 3112113 | 180 | D | Conv. |
| CS05 | Gran Canaria | 35/6/8/272/1 | 28°08'24'' N | 15°31'03'' W | 28 R 449184 3112819 | 135 | D | Conv. |
| CS06 | Gran Canaria | 35/6/8/455/3 | 28°08'10'' N | 15°31'23'' W | 28 R 448636 3112391 | 135 | D | Conv. |
| CS07 | Gran Canaria | 35/9/1/244/1 | 28°09'11'' N | 15°39'58'' W | 28 R 434598 3114337 | 17 | D | Conv. |
| CS08 | Gran Canaria | 35/6/7/312/1 | 28°08'14'' N | 15°31'36'' W | 28 R 448282 3112515 | 135 | D | Conv. |
| CS09 | Gran Canaria | 35/9/1/260/4 | 28°08'49'' N | 15°40'07'' W | 28 R 434349 3113661 | 32 | D | Conv. |
| CS10 | Gran Canaria | 35/9/1/288/1 | 28°09'03'' N | 15°39'52'' W | 28 R 434761 3114090 | 40 | D | Conv. |

**Table S2. Fungal volatile organic compounds selected for the olfactometry bioassays.**

| **Num.** | **Compound** | **Ret. Time (min)** | **Peak Height 10d** | **Peak Height 20d** | **Peak Height 30d** | **Peak Height 40d** | **Peak Height 50d** | **Peak Height 60d** | **Fungal species** |
| --- | --- | --- | --- | --- | --- | --- | --- | --- | --- |
| **C1** | **Styrene** | 18.55 | **-** | **-** | **-** | **-** | **-** | **-** | Bb1TS11 Bb203 |
| **C2** | **Benzothiazole** | 31,00 | NA | NA | 23152 | NA | NA | NA | Bb1TS11  (Bb203) |
| **C3** | **Camphor** | 29.11 | **-** | **-** | **-** | **-** | **-** | **-** | Bb1TS11 |
| **C4** | **Borneol** | 29,30 | 96273 | 24965 | NA | 31486 | 20551 | NA | Bb1TS11 |
| **C4** | **Borneol** | 29,30 | 52467 | 48687 | 39535 | 39445 | 66449 | 40861 | Bb203 |
| **C5** | **1,3-dimethoxy-benzene** | 28,75 | 263989 | NA | NA | NA | NA | NA | Ma4TS04 |
| **C5** | **1,3-dimethoxy-benzene** | 28,75 | 7233098 | 509769 | 3583189 | 3807170 | 3046943 | 308330 | Pc123 |
| **C6** | **1-octen-3-ol** | 22,36 | 26983 | 385375 | 141833 | 28896 | NA | NA | Ma4TS04 |
| **C6** | **1-octen-3-ol** | 22,36 | 123365 | 108363 | 557934 | NA | NA | 308377 | Pc123 |
| **C7** | **3-cyclohepten-1-one** | 15,20 | 50086 | 475717 | 306136 | 120587 | NA | 233415 | Bb1TS11 |
| **C7** | **3-cyclohepten-1-one** | 15,20 | 274698 | 246124 | 103529 | NA | NA | NA | Ma4TS04 |
| **C7** | **3-cyclohepten-1-one** | 15,20 | 714704 | 1027453 | NA | NA | NA | NA | Pc123 |
| **C7** | **3-cyclohepten-1-one** | 15,20 | 179843 | 704724 | 147859 | 330871 | 239657 | 263098 | Bb203 |

**Table S3. Volatile organic compounds produced by the fungi analysed, in all the times measured.** The list is made with a selection of compounds with a match percentage on the databases of at least 50%, without limitations in abundance.

| Compound number | Compound name | Retention time (min) | Fungi |
| --- | --- | --- | --- |
| 1 | 2-butanone | 6,95 | Pc123 |
| 2 | Methyl ethyl, 2-butanone | 6,99 | Pc123 |
| 3 | 3-methylbutanal | 9,10 | Pc123 |
| 4 | 2-methylbutanal | 9,43 | Pc123 |
| 5 | Dimethyldisulfide | 12,60 | Pc123 |
| 6 | 3-hydroxy-2-butanone | 12,76 | Pc123 |
| 7 | 4-methyl-2-pentanone | 13,04 | Pc123 |
| 8 | 3-methyl-1-butanol | 13,28 | Ma4TS04 |
| 9 | 2-chloro-octane | 13,45 | Ma4TS04 |
| 10 | 3-methyl-2-pentanone | 13,46 | Pc123 |
| 11 | Cis-1-butyl-2-methylcyclopropane | 13,49 | Bb1TS11 |
| 12 | 1-octene | 13,50 | Bb203; Bb1TS11 |
| 13 | 1,3-octadiene | 15,20 | Ma4TS04; Bb1TS11 |
| 14 | 3-cyclohepten-1-one | 15,20 | Ma4TS04; Pc123; Bb203; Bb1TS11 |
| 15 | Methyl-1,4-dioxide-pyrazine | 15,24 | Bb1TS11 |
| 16 | (1-methylethenyl) cyclopropene | 15,31 | Pc123 |
| 17 | 3-hydroxy-2-pentanone | 16,41 | Pc123 |
| 18 | Diethylamine-D1 | 16,41 | Pc123 |
| 19 | 2,4-octadiene | 16,64 | Pc123; Bb203 |
| 20 | 4-methyl-3-hexanone | 16,90 | Pc123 |
| 21 | Nonane | 17,17 | Pc123 |
| 22 | 3-methyl-3-penten-2-one | 17,18 | Pc123 |
| 23 | 5-methyl-2-hexanone | 17,83 | Pc123 |
| 24 | α-pinene | 19,40 | Pc123; Bb203 |
| 25 | 2-butoxy-ethanol | 19,79 | Pc123 |
| 26 | Anisole | 19,80 | Pc123 |
| 27 | Methoxybenzene | 19,82 | Pc123 |
| 28 | Propionoin | 20,01 | Pc123 |
| 29 | 2,4-dimethyl-3-hexene | 20,54 | Pc123 |
| 30 | 3,4-dimethyl-2-hexene | 20,55 | Pc123 |
| 31 | Trans-3,4-dimethyl-2-hexene | 20,55 | Pc123 |
| 32 | 3,5-dimethyl-2-propylthiophene | 20,61 | Ma4TS04 |
| 33 | β-pinene | 21,20 | Pc123; Bb203 |
| 34 | 1-β-pinene | 21,24 | Ma4TS04 |
| 35 | 2,2,4,6,6-pentamethyl-heptane | 21,31 | Bb1TS11 |
| 36 | 2-isopropyl-5-oxohexanal | 21,42 | Pc123 |
| 37 | 6-methyl-2-heptanone | 21,42-25,13 | Pc123 |
| 38 | Propyl-cyclohexane | 21,69 | Pc123 |
| 39 | Dimethyl-trisulfide | 21,79 | Pc123 |
| 40 | (2R*,6R*,8AS*)-6-hydroxyedulan | 22,01 | Bb203 |
| 41 | 2-methyl-propanamide | 22,14 | Pc123 |
| 42 | 1-octen-3-ol | 22,36 | Ma4TS04; Pc123 |
| 43 | 3-octanone | 22,40 | Pc123; Bb203 |
| 44 | 3-methoxy-benzenamine | 22,56 | Ma4TS04 |
| 45 | Trimethyl-pyrazine | 22,70 | Pc123 |
| 46 | 2-methyl-2-bornene | 22,71 | Pc123; Bb203; Bb1TS11 |
| 47 | 2,4-dimethyl-methyl ester-hexanoic acid | 22,95 | Pc123 |
| 48 | 6-methyl-bicyclo[3.3.0]oct-2-en-7-one | 23,04 | Bb203 |
| 49 | 1-methyl-4-(1-methylethyl)-benzene | 23,18 | Ma4TS04; Pc123 |
| 50 | Isocyano-benzene | 23,58 | Pc123 |
| 51 | 2-(2-ethoxyethoxy)-ethanol | 23,60 | Ma4TS04; Pc123; Bb203; Bb1TS11 |
| 52 | 2,2,4,4,6,8,8-heptamethyl-nonane | 24,26 | Bb1TS11 |
| 53 | 2,4-dimethylfuran | 24,41 | Pc123 |
| 54 | 1,1'-oxybis-heptane | 24,77 | Bb203 |
| 55 | 2-dodecanone | 24,9-34,28 | Pc123 |
| 56 | 2-methyl-2-(2-methyl-2-butenyl)-furan | 25,47 | Pc123 |
| 57 | Benzenemethanol | 25,49 | Pc123 |
| 58 | (2-methyl-1-propenyl)-benzene | 25,57 | Ma4TS04 |
| 59 | 1-methyl-1H-1,2,4-triazole | 25,58 | Bb1TS11 |
| 60 | 2,3,3-trimethyl-1-butene | 25,58 | Bb203 |
| 61 | 2,2-dimethyl-propanal | 25,63 | Ma4TS04 |
| 62 | 2-methyl-3,4-dihydro-2H-pyran | 25,70 | Bb1TS11 |
| 63 | 6-methyl-4 5-dihydro.alpha.[2H]-pyran | 25,70 | Pc123 |
| 64 | 6,7-dimethoxy-2,2-dimethyl-2H-1-benzopyran | 26,41 | Pc123 |
| 65 | 2-amino-3,5-dibromo-6-methylpyridine | 26,49 | Ma4TS04 |
| 66 | 3-hydroxymandelic acid ethyl ester di-TMS | 26,49 | Pc123 |
| 67 | 4-hydroxymandelic acid ethyl ester di-TMS | 26,49 | Ma4TS04; Pc123; Bb203 |
| 68 | 4-trimethylsilyl-9,9-dimethyl-9-silafluorene | 26,49 | Pc123 |
| 69 | p-Trimethylsilyloxyphenyl-bis(trimethylsilyloxy)ethane | 26,50 | Bb203 |
| 70 | 1-methylallyl(cyclooctatetraene)titanium | 27,22 | Pc123; Bb1TS11 |
| 71 | 4-octen-3-ol | 27,55 | Ma4TS04 |
| 72 | 4-fluoro-1,2-xylene | 27,90 | Ma4TS04; Bb203 |
| 73 | 1-phenyl-2-propanone | 28,22 | Pc123 |
| 74 | 1,3-dimethoxy-benzene | 28,75 | Ma4TS04; Pc123 |
| 75 | 2-amino-4-methylpyrimidine | 29,14 | Pc123 |
| 76 | Bicyclo[2.2.1]heptane-2-carboxylic acid | 29,14 | Pc123 |
| 77 | Borneol | 29,30 | Bb203; Bb1TS11 |
| 78 | 9H-pyrrolo[3',4':3,4]pyrrolo[2,1-a] phthalazine-9,11(10H)-dione,10-ethyl-8-phenyl | 30,54 | Ma4TS04 |
| 79 | 1-(2-furanyl)-3-butene-1,2-diol | 30,96 | Ma4TS04 |
| 80 | 2,4-bis(1,1-dimethylethyl)-phenol | 31,00 | Ma4TS04; Pc123 |
| 81 | Benzothiazole | 31,00 | Bb1TS11 |
| 82 | 2-undecanone | 31,15-32,18 | Pc123 |
| 83 | 16-oxosalutaridine | 31,48 | Pc123 |
| 84 | 5-oxoisoboldine | 31,48 | Ma4TS04 |
| 85 | 4,4-dimethyl-6-benzylamino-dihydro-8H-thiopyrano[4',3':4,5]thieno[2,3-d]pyrimidine | 31,50 | Ma4TS04; Pc123 |
| 86 | 2-octanone | 32,19 | Pc123 |
| 87 | 1-ethyl-2-methyl-cyclohexane | 33,21 | Pc123 |
| 88 | 5-ethyl-3-hydroxy-4-methyl-2(5H)-furanone | 33,21 | Bb1TS11 |
| 89 | 5-acetyl-2-hydrazino-4-methylpyridine | 33,70 | Pc123 |
| 90 | β-elemene | 34,61 | Pc123 |
| 91 | 2-methoxy-3,8-dioxocephalotax-1-ene | 34,86 | Pc123 |
| 92 | Di-amylcyclohexanol | 34,97 | Pc123 |
| 93 | Cyclotetradecane | 36,68 | Pc123 |
| 94 | (+-)-gymnomitrene | 37,10 | Ma4TS04; Bb1TS11 |
| 95 | 1,4-bis(methylene)-cyclohexane | 37,10 | Ma4TS04 |
| 96 | 2-tridecanone | 37,60 | Pc123 |
| 97 | trans-β-farnesene | 37,94 | Pc123 |
